# Spatial whitening in the retina may be necessary for V1 to learn a sparse representation of natural scenes

**DOI:** 10.1101/776799

**Authors:** Eric McVoy Dodds, Jesse Alexander Livezey, Michael Robert DeWeese

## Abstract

Retinal ganglion cell outputs are less correlated across space than are natural scenes, and it has been suggested that this decorrelation is performed in the retina in order to improve efficiency and to benefit processing later in the visual system. However, sparse coding, a successful computational model of primary visual cortex, is achievable under some conditions with highly correlated inputs: most sparse coding algorithms learn the well-known sparse features of natural images and can output sparse, high-fidelity codes with or without a preceding decorrelation stage of processing. We propose that sparse coding with biologically plausible local learning rules does require decorrelated inputs, providing a possible explanation for why whitening may be necessary early in the visual system.

## 1 Introduction

Following proposals that the brain seeks to reduce redundancy in signals from the natural environment[2, 3], such as natural visual scenes [17, 9], Atick and Redlich proposed that the center-surround receptive fields of retinal ganglion cells serve to decorrelate natural visual input to obtain a representation with less redundancy among the outputs of individual neurons, while also suppressing noise [1]. Atick and Redlich emphasized advantages of a code that uses statistically independent elements, such as simple computation of joint probabilities. The removal of pairwise linear correlations, or “whitening”, is then seen as a first step towards such a representation. In fact, there is strong experimental evidence for decorrelation at the earliest stages of the visual pathway [7, 11], including nonlinear retinal processing to remove spatial correlations [21]. Studies have also suggested that fixational eye movements may contribute to decorrelation of retinal ganglion cell outputs[25, 16].

In this paper, we propose that it may in fact be *necessary* to decorrelate visual input before circuitry in primary visual cortex (V1) can achieve a sparse representation of natural scenes if plasticity mechanisms at each synapse can only access local information.

A popular normative theory for V1 simple cells postulates that these neurons are optimized for forming sparse representations of natural visual stimuli [10, 20]. Olshausen and Field demonstrated that a sparse coding model trained on photographs can learn visual features that resemble receptive fields measured in primate V1 simple cells [20]. Related algorithms obtain similar results [4] or even closer agreement with experiment [23, 29]. Similar models have also been successfully applied to other sensory modalities [26, 15, 6].

The “Sparse and Independent Local Network” (SAILnet) [29], a sparse coding network with more biologically realistic spiking neurons and local learning rules, obtains similarly strong results when trained on whitened natural image patches. A modified version of SAILnet with separate populations of excitatory and inhibitory neurons also produces reasonable receptive field shapes [14].

Whereas sparse coding studies often use whitening as a preprocessing step, pre-whitening is not necessary for conventional methods, such as those of [20, 4, 23]. In fact, some authors have pointed to the existence of nonlinearities in the retina and thalamus as an argument against sparseness as a normative theory for coding in V1 [19]. We show that SAILnet, however, does not learn V1-like receptive fields if the image data is not pre-whitened. SAILnet differs from conventional sparse coding in several ways, but the essential feature that makes pre-whitening necessary is the network’s synaptically local learning rules.

We call a learning rule “synaptically local” if the strength of a synapse is updated using information known to be available at that synapse: the number and timing of recent spikes in the pre-synaptic and post-synaptic neurons and the present strength of the synapse.

Importantly, sparse coding models with *non*-local plasticity can perform well on unwhitened data such as raw natural images. We demonstrate this with an algorithm that finds sparse codes using the Locally Competitive Algorithm (LCA) [24] to perform inference and stochastic gradient descent (SGD) on mean squared error, a non-local learning rule, to perform learning. While this network learns somewhat more accurately and quickly on whitened data than on unwhitened data, pre-whitening is not necessary to generate codes of high fidelity and sparseness or to learn visual features resembling V1 receptive fields.

To summarize, this paper makes two contributions: (1) we propose that biologically plausible local learning rules may require decorrelated inputs to learn a sparse model of natural scenes in V1, a novel possible explanation for whitening occurring in a separate, earlier stage of the visual system; and (2) we demonstrate that conventional nonlocal learning rules do not require whitened inputs, while providing evidence that biologically plausible local learning rules do require whitened inputs. This evidence takes the form of general arguments plus a demonstration in the important specific case of SAILnet.

## 2 Results and Discussion

We demonstrated the contrast between learning with SAILnet versus conventional sparse coding on whitened and unwhitened data in two complementary ways, both shown in Fig. 1. First, we trained models on raw natural image patches and also on the same patches after whitening. We trained SAILnet with many different settings of its hyperparameters; results corresponding to the optimal hyperparameters (*i.e.*, those that give the highest-fidelity sparse codes) are shown in the figure. No values of these hyperparameters led to SAILnet learning qualitatively different receptive fields in the unwhitened case. In particular, we never observed SAILnet to learn receptive fields with localized structure from unwhitened natural images. Rather, SAILnet always learned features like those in Fig. 1D, with the lowest spatial frequencies dominating. Components with low spatial frequencies account for the greatest proportion of the variance of (unwhitened) natural images [9] but do not allow the network to efficiently represent finer-grained detail.

**Figure 1:**
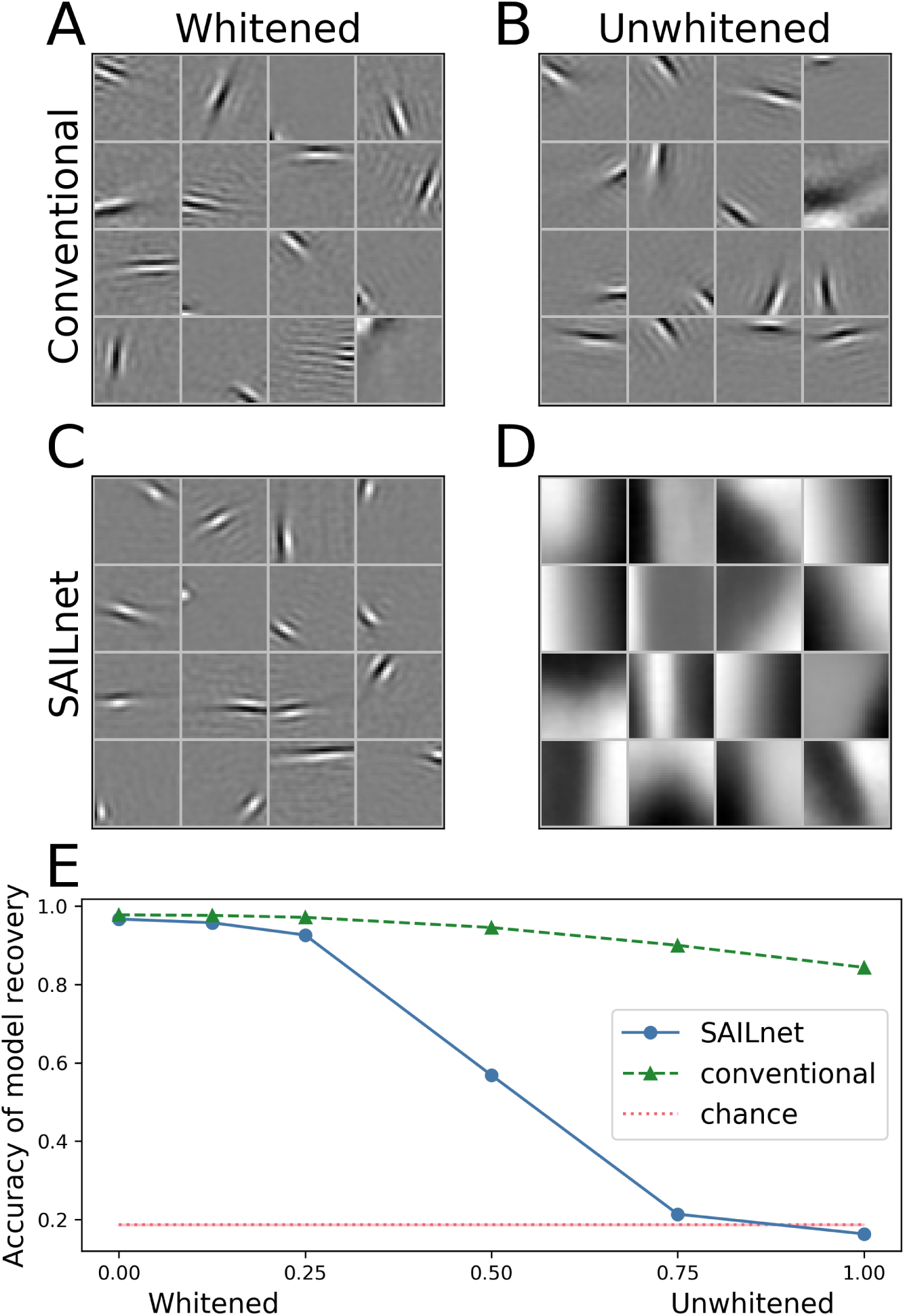
SAILnet learning requires pre-whitening while conventional non-local learning does not. (A) Receptive fields of conventional sparse coding model neurons resemble V1 simple cell receptive fields after the network has been trained on a set of whitened grayscale natural image patches. The network had 256 units; a random sample of 16 units are shown. (B) Conventional sparse coding learns qualitatively similar features on raw, unwhitened natural image patches. (C) SAILnet receptive fields after training on a set of whitened natural image patches resemble conventional results and V1 simple cells. (D) SAILnet receptive fields after training on unwhitened image patches are markedly different. (E) Results for SAILnet and conventional sparse coding on a sparse model recovery task where the data has been de-whitened to some degree, with 1 on the horizontal axis corresponding to the same pairwise correlation structure as natural images. Whereas conventional methods degrade slightly with de-whitening, SAILnet performs no better than a randomly generated dictionary (dotted red line) for synthetic data as far from white as natural images. All simulations were run five times. Means are plotted; standard deviations are too small to display as error bars.

Second, we generated synthetic data consisting of sparse (Laplace-distributed) linear combinations of a fixed, randomly-generated set of vectors; each of these vectors was an instantiation of frozen white noise. Both conventional sparse coding and SAILnet were able to recover these vectors from the data. We then applied to this synthetic data the singular value spectrum of our natural image patch dataset — in effect, “unwhitening” the dataset. We trained models on this type of synthetic data with distributions of singular values interpolated between that of natural images and the original whitened data, searching many values of SAILnet hyperparameters in each case. We found that conventional sparse coding continued to perform well for all spectra we tested, whereas SAILnet’s performance collapsed to chance-level for data with the singular value spectrum of natural images (Fig. 1E). Just as the network learned only features dominated by the high-variance, low-spatial-frequency components of natural images when trained on unwhitened natural images, the network learned features dominated by whichever components have the highest variance in the synthetic data. See Sec. 3.4 for more details on this experiment.

We have been unable to find, either in the literature or by our own efforts, a sparse coding algorithm using synaptically local learning rules of the type discussed above that performs well on unwhitened natural images. We believe that any such algorithm optimizing for sparse coding will tend to be pulled towards directions of high variance (in the case of natural images, features dominated by low spatial frequencies) just as SAILnet is.

More specifically, a network with local learning rules needs to overcome two related challenges: 1) learning lateral connections to facilitate cooperation in coding without each neuron having direct access to the stimulus features that other neurons represent; and 2) learning feedforward connections to minimize future coding errors without each neuron having direct access to the contribution to the representation of the stimulus by other neurons. SAILnet’s inhibitory connection learning rule solves the first problem by leveraging the fact that neuron activity correlations are closely tied to similarity of the features the neurons represent, but strong correlations in the stimulus can distort this relationship. Feedforward synapses onto a given neuron lack direct access to other neurons’ spikes, so they must learn to reduce coding error using only the stimulus and the neuron’s spikes. If some directions in stimulus space have higher variance than others, these synapses can best reduce error alone within the spike budget imposed by the sparseness constraint by aligning with the high-variance directions, overwhelming the advantage of a new stimulus direction of sufficiently lower variance.

Following work on biologically plausible coding with excitatory-inhibitory balance [5][8], we also developed sparse coding learning rules for which each synapse also has access to a neuron-wide “membrane potential” variable that represents the summed synaptic input to the neuron. We find that these rules can learn sparse representations from unwhitened data (see Sec.4.2). However, dendrites in real neurons are typically electrically compartmentalized [13, 22, 12], making it unlikely that the membrane potential as measured in the soma is universally available across all synapses throughout the dendritic tree of any given neuron. In principle, a synapse could rely on a known relationship between the local membrane potential and that of the soma, but changes in synaptic strengths throughout the dendritic tree would alter that relationship during learning. Moreover, especially at distal synapses, the membrane potential at any given point in the dendrite would not be deterministically related to the somatic membrane potential due to fluctuations in the input and channel noise, even for a fixed set of synaptic weights throughout the dendritic tree.

It has also been shown that a sparse coding network can learn successfully from unwhitened inputs by comparing sparse codes for the same stimulus generated by two different configurations of the network [18]. Whereas the learning rules in this method use information local to each synapse, we do not think it is biologically plausible that each synapse could acquire, store, and compare the spike rates for two completely different network configurations at each moment in time.

While we believe it is unlikely, more complicated models that make use of precise relative spike timing could in principle obviate the need for pre-whitening while using local learning rules to learn a sparse code. It is not clear how such a scheme could overcome the challenges described above.

Our proposal compliments the prevailing notion that retinal whitening provides a more efficient representation for transmission through the limited capacity of the optic nerve before the creation of an overcomplete, sparse representation in primary visual cortex, and it suggests a second compelling reason for decorrelation to be performed by a distinct population of neurons at the earliest stages of the visual system.

## 3 Methods

### 3.1 Data and whitening

We used a set of *N* = 3 ×10^5^ square patches of 256 pixels each drawn from the van Hateren image dataset[28]. The full images, but not the patches, were mean-centered and normalized by the standard deviation across pixels. We then subtracted the mean patch and divided by the standard deviation across all pixels and patches.

Whitening or sphering refers to a linear transformation that results in a covariance matrix proportional to the identity. In this work we use PCA whitening. First we multiply the data by the matrix of eigenvectors of the covariance matrix, and then we divide each component by its singular value. So the whitened dataset of *N* examples, *X* ∈ ℝ^*N*×*M*^, is

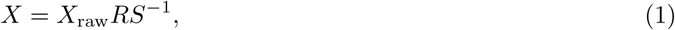

where *S* is an *M* × *M* diagonal matrix containing the singular values of *X*_raw_, *R* ∈ *O*(*M*) is a rotation, and

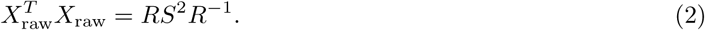

A 2D visualization is shown in Fig. 2.

**Figure 2:**
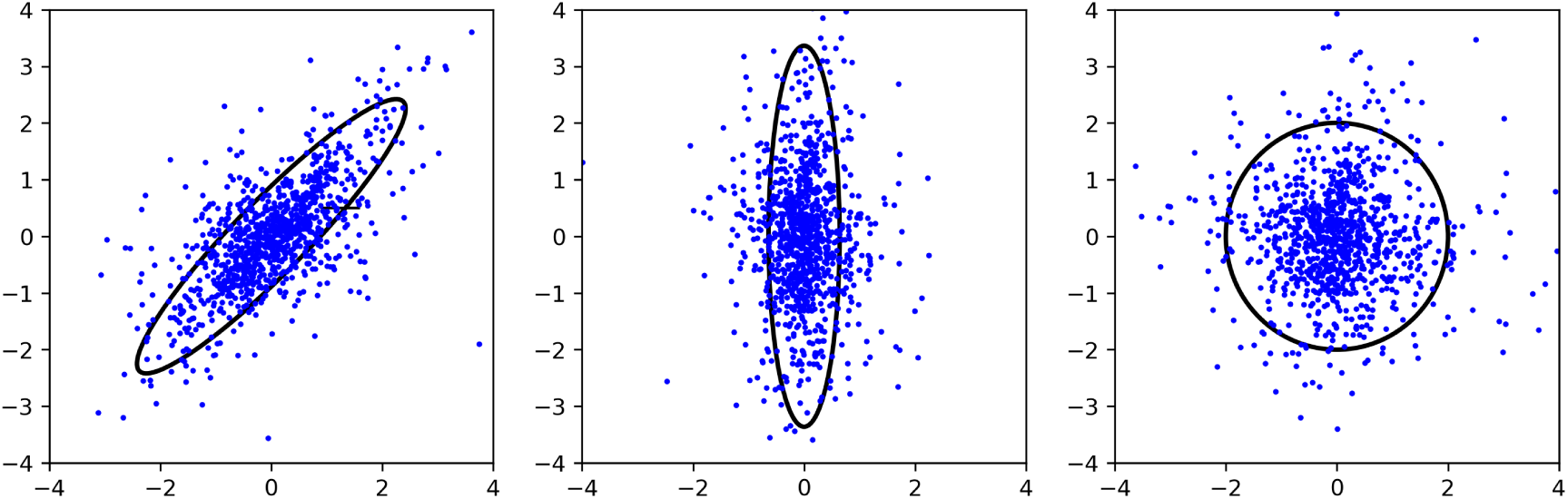
PCA whitening for 2D data. Data can be whitened by transforming to the principal component basis and then scaling the axes (that is, the principal components). The result and all rotations of the result are whitened.

The distribution of standard deviations of the principal components, also known as the singular values of the data, characterizes the asphericity of the data. In the case of natural images (and other data with approximate translational invariance), the singular values correspond closely with the Fourier amplitude spectrum.

### 3.2 Sparse coding

We first define a probabilistic model

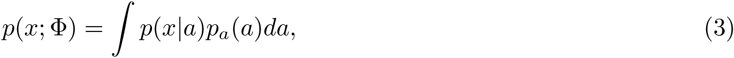

where *x* denotes a data vector with components *x*_*i*_ and *a* denotes a set of latent variables *a*_*m*_. The conditional distribution *p*(*x*|*a*) is an isotropic Gaussian of fixed variance centered on a linear reconstruction of the data in terms of the dictionary elements Φ_*m*_:

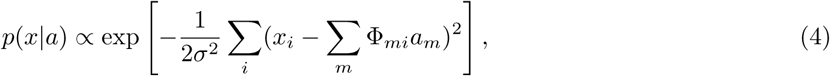

and the prior distribution of the coefficients *a*_*m*_ is factorial with each factor given by the same sparse distribution (*e.g.*, a Laplace distribution):

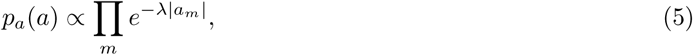

where *λ* is a parameter that determines the width of the distribution and therefore how strongly the prior favors sparse sets of *a*_*m*_.

Fitting this model usually involves estimating the gradient of the likelihood (or log-likelihood) with respect to the parameters Φ_*mi*_, but this calculation involves an intractable integral over the latent variables *a*_*m*_. A common approximation is to set the *a*_*m*_ by maximum *a posteriori* (MAP) inference given input data *x*:

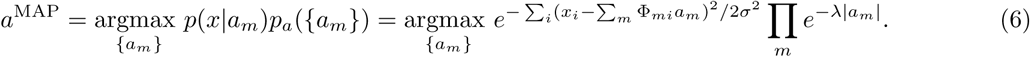

The parameters *σ*^2^ and *λ* now only affect the model through the combination *λσ*^2^, so to simplify the notation we set *σ* = 1.

The quantity 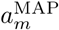 is typically referred to as the activity of the *m*th unit, and the dictionary elements {Φ_*m*_} are often compared to receptive fields of neurons. The analogies to neurons suggested by these terms are not exact, but a unit’s dictionary element is approximately the same as the linear receptive field that would be measured for that unit with an activity-triggered average [20].

The dictionary elements Φ_*m*_ are conventionally learned by descending the estimate of the gradient provided by differentiating the model log-likelihood with respect to Φ with *a* fixed at the MAP value:

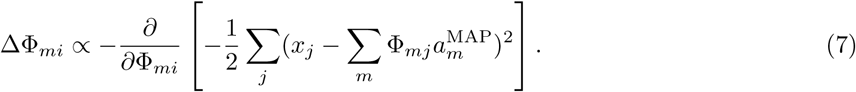

In this work, this gradient with respect to Φ is averaged over a minibatch of 100 data examples.

The use of MAP inference requires that we constrain the norms of the Φ_*m*_ to avoid solutions with small *a*_*m*_ and large, meaningless Φ_*m*_. We therefore divide each Φ_*m*_ by its norm after each gradient step.

#### 3.2.1 Locally Competitive Algorithm

We use the L1-sparse locally competitive algorithm (LCA) [24] to perform MAP inference in our “conventional sparse coding” model. LCA uses a dynamical system with auxiliary variables that are thresholded to obtain estimates of *a*^MAP^. Typically most of the auxiliary variables are below threshold and the 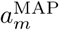 estimates are exactly zero for most *m*. The threshold is set by the sparseness parameter *λ*.

The choice of coding algorithm is not crucial to our results, and learning using alternative inference schemes yields qualitatively similar dictionaries.

### 3.3 SAILnet and related models

SAILnet uses leaky integrate-and-fire neurons to form a sparse spike-count code and local rules to updates its synaptic strengths. The structure of the network is illustrated in Fig. 3.

**Figure 3:**
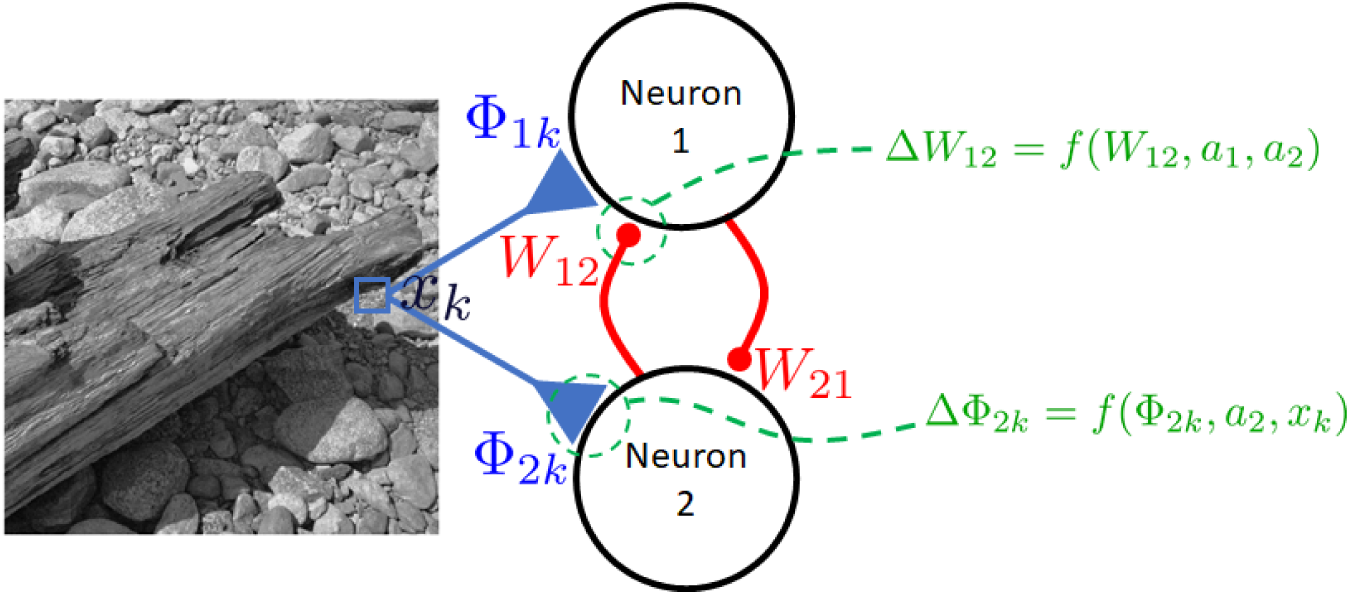
SAILnet and similar models use feedforward and lateral connections updated with synaptically local learning rules. Changes in the strength of each synapse depend only on quantities available at that synapse, including the current strength of that synapse and post- and pre-synaptic spikes.

The LIF unit dynamical equation is

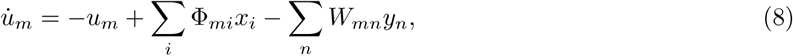

where the spike variable *y*_*n*_ is 0 unless the membrane potential *u*_*m*_ just crossed the threshold *θ*_*m*_ (after each spike, *u*_*m*_ is reset to 0). (Here and elsewhere we use time units such that the resulting time constant is 1.) An example LIF unit’s evolution is shown in Fig. 4. (Replacing the LIF neurons with continuous-valued units similar to LCA units makes a quantitative but not qualitative difference to learning.) These dynamics give approximately optimal codes *a*_*m*_, proportional to the spike counts, if the lateral weights are 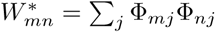

**Figure 4:**
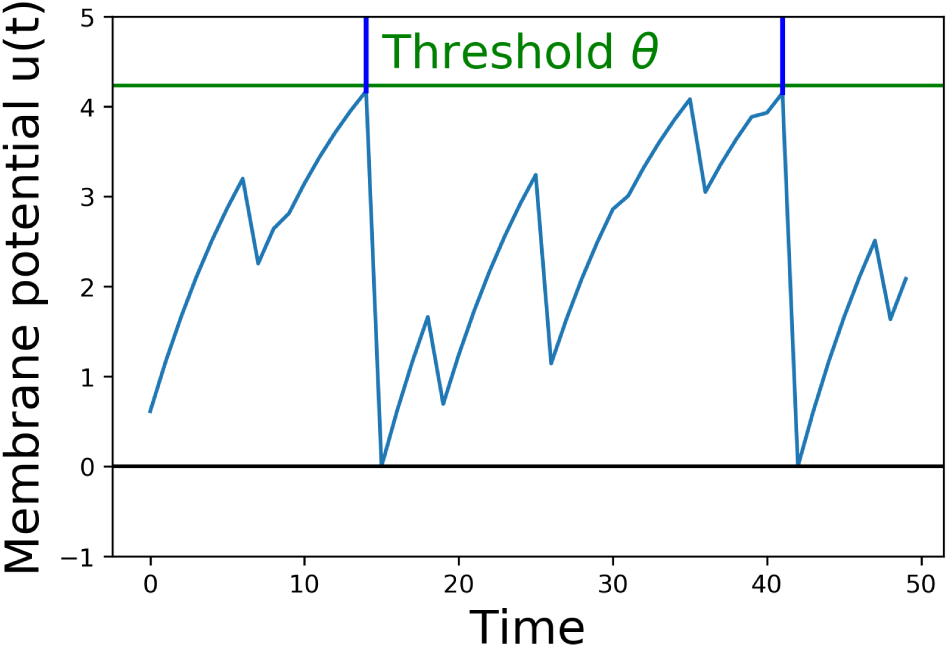
SAILnet uses leaky integrate-and-fire (LIF) model neurons. Constant feedforward inputs from the data drive the unit’s internal “membrane potential” variable *u*, while inhibitory inputs from other units decrease *u* whenever the inhibiting units spike. When *u* hits the unit’s threshold *θ*, the unit emits a spike and resets *u* to zero at the next timestep. Each unit has its own learned threshold *θ*.

SAILnet updates its parameters according to

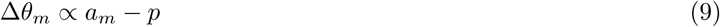

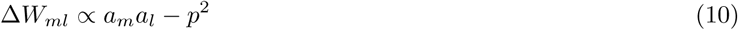

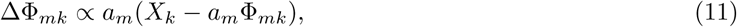

where *a*_*m*_ is the (possibly normalized) spike count for unit *m*. The first equation imposes a constraint on the firing rate of each neuron. The second attempts to impose the constraint that the neurons’ activities be decorrelated. The third drives each neuron to reconstruct the stimulus (alone).

### 3.4 Synthetic data

Our synthetic data was generated by sampling from the sparse coding probabilistic model (3) with a known, fixed dictionary. For *N* sources in *D* dimensions, we generated the dictionary Φ^*^ by sampling *N* directions in *D*-dimensional space uniformly at random. Then each data vector *x*^*µ*^ was determined by *N* samples 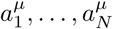 from an exponential distribution with scale *λ*:

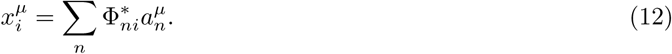

This data generation process is spherically symmetric, but any particular dictionary and dataset will not be perfectly white. We used *D* = 256 and *N* = 256 for our experiments, for as close correspondence with the natural image experiments as possible.

To de-whiten the synthetic data, we multiplied by the matrix of singular values of our natural image patch dataset, raised to some power *ψ*:

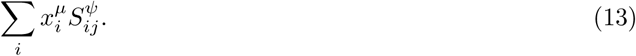

The parameter *ψ* is represented by the horizontal axis of Fig. 1E.

To quantify the quality of the model recovery, we calculated the largest dot product of a learned dictionary element for each ground-truth dictionary element:

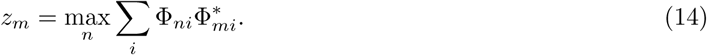

Recall that all *ψ*_*n*_ and *ψ* _\**n*_ are normalized. The median value of the *z*_*m*_ across dictionary elements is represented by the vertical axis of Fig. 1E.

Models were trained on the synthetic data starting from random initial dictionaries Φ. The elements of these random dictionaries are not in general orthogonal to the target dictionary elements, so our model recovery measure has a non-zero “chance” value which we estimated and plot as the red line in Fig. 1.

## 4 Supporting information

### 4.1 Complete dictionaries of learned models

We believe the sample dictionary elements in Fig. 1 represent the learned dictionaries faithfully for the purposes of our argument, but we present here the complete dictionaries (Figs. S1, S2, S3, and S4). In each case the dictionary elements, which are trained on principal component representations, are projected back to image space for presentation with no other alteration. In particular, the whitening matrix is not used in this transformation.

### 4.2 Models using membrane potential and spike times

There are other notions of locality and “biological plausibility” that do not fit within the framework described in the main text. One possibility is to allow learning based on information that may be available during the “inference” process but that isn’t in the final activations *a*_*m*_. Inspired by the work of [5, 8] on “learning to represent signals spike by spike,” we developed a SAILnet-like sparse coding model in which the post-synaptic membrane potential is used in addition to spikes to determine synaptic strength updates. This model, which we describe below, learns the expected Gabor-like features on natural image patches regardless of whether the data is pre-whitened, and recovers a ground-truth model when the data has been de-whitened (Fig S5).

#### 4.2.1 Sparse coding spike by spike

In the supplementary information of [5], Brendel et al. provide a set of learning rules for an efficient autoencoding network that can be trained on unwhitened data. In our notation and framework the feedforward weight rule is

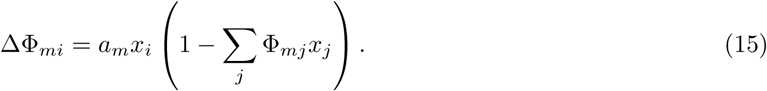

A modified version of SAILnet incorporating this rule and the membrane potential-based *W* rule can learn using unwhitened data.

We emphasize that this rule as written does not fit our notion of biological plausibility: each synapse needs to know an aggregate of many other synapses’ inputs that is not simply captured in the membrane potential.

**Figure S1:**
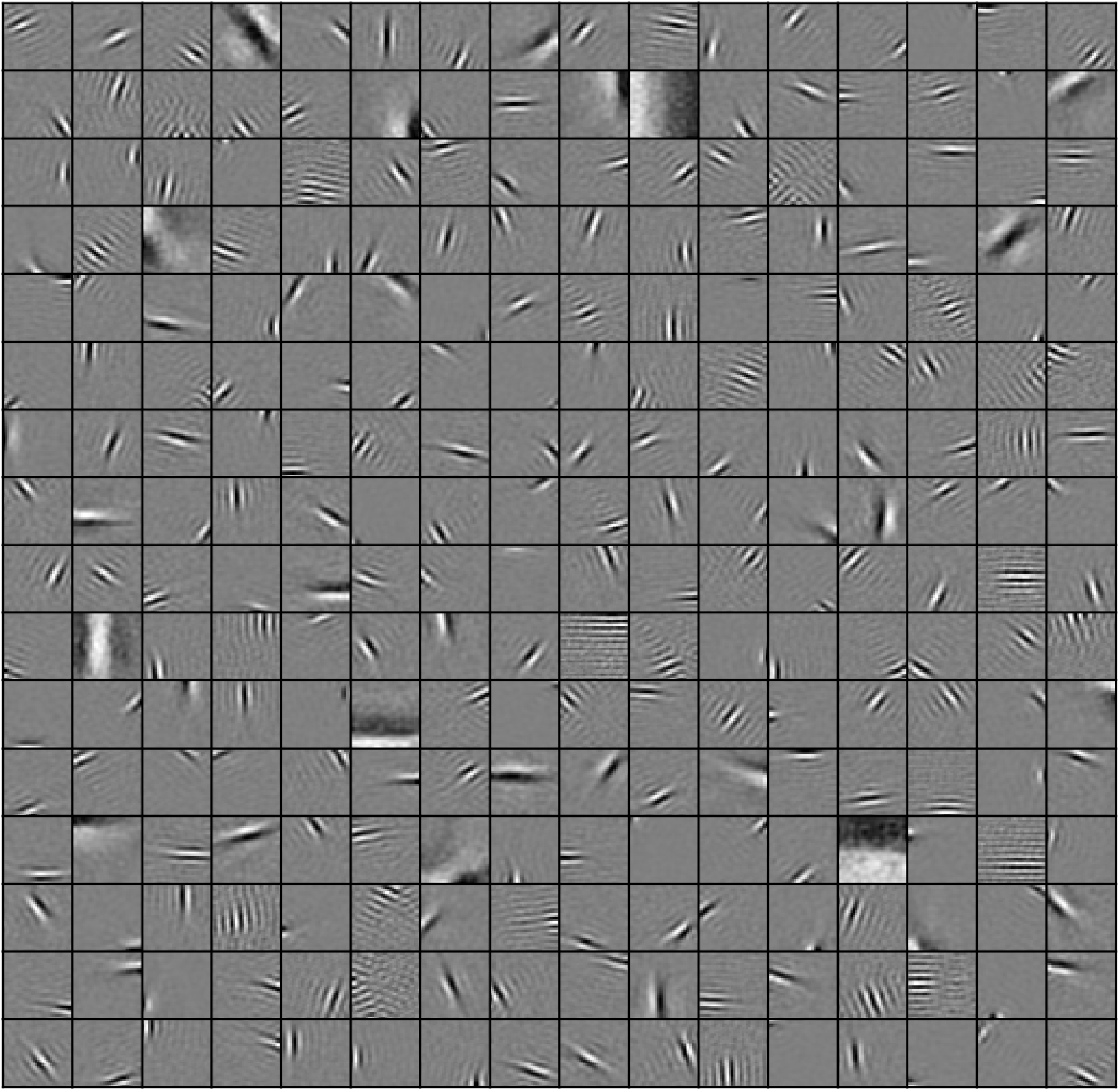
All dictionary elements from a complete sparse coding dictionary learned with stochastic gradient descent and LCA inference on whitened natural image patches.

However, an approximation using the membrane potential in place of this term also works:

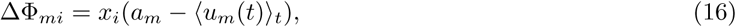

where the angle brackets denote averaging over inference time. To achieve sparse codes, we can use this learning rule together with the inhibitory weight learning rule of [5] and the SAILnet spiking threshold learning rule of [29]. For completeness, the inhibitory weight learning rule in our notation is

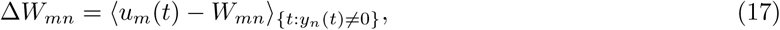

where the angle brackets denote an average over the post-synaptic neuron’s spike times. We call the resulting network MP-SAILnet, using the rules in Eqs. 16 and 17 plus the SAILnet threshold update. MP-SAILnet can learn the expected sparse features of natural image patches without pre-whitening, at the cost of requiring a neuron-global “membrane potential” to be available to the feedforward synapses while determining their weight updates. We discuss this issue further and clarify the comparison to SAILnet and other models in section 4.3.

**Figure S2:**
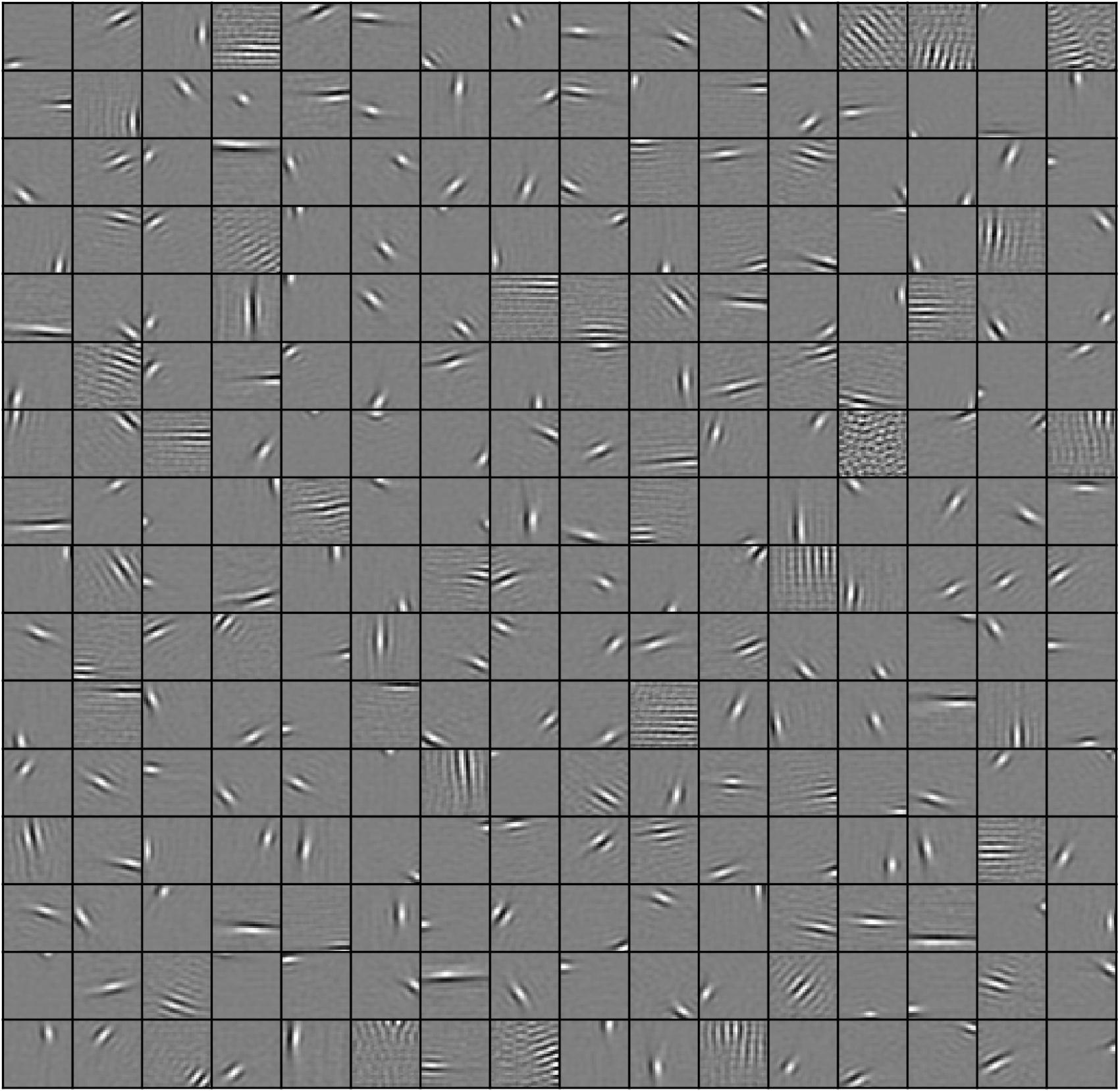
All dictionary elements from a complete SAILnet model trained on whitened natural image patches.

### 4.3 Model variations and relation to whitening

A number of neurally-inspired sparse coding models have been proposed to learn the parameters of sparse linear generative models. The properties of these models are “biologically plausible” to different degrees and in different ways, and the various algorithms have various strengths and weaknesses in perfomance. Here we attempt to clarify the type of biological plausibility we have focused on in this work and the ways in which models that are more or less plausible in this sense can or cannot learn effectively from data that is not pre-whitened.

The primary aspect of biological plausibility we are concerned with here is the *locality* of the learning rules. Locality can mean several things with regard to the neural circuit. For example, one possibility is that learning rules should be “neuron-local”, meaning that they should only depend on quantities that at least some part of a neuron would have access to. For instance, this would include: all of the synaptic strengths in the given neuron’s dendritic tree, Φ_*mj*_, where *j* labels the presynaptic input and *m* is the index labeling the neuron; the neuron’s somatic membrane potential and spiking output, *u*_*m*_ and *a*_*m*_, respectively; and the spiking activity of each of the presynaptic neurons, *a*_\*m*_, where the \*m* index refers to all neurons other than the postsynaptic neuron. By contrast, it is unlikely that any part of the neuron would have access to the synaptic strengths in the dendrites of other neurons: Φ_(\*m*)*j*_.

**Figure S3:**
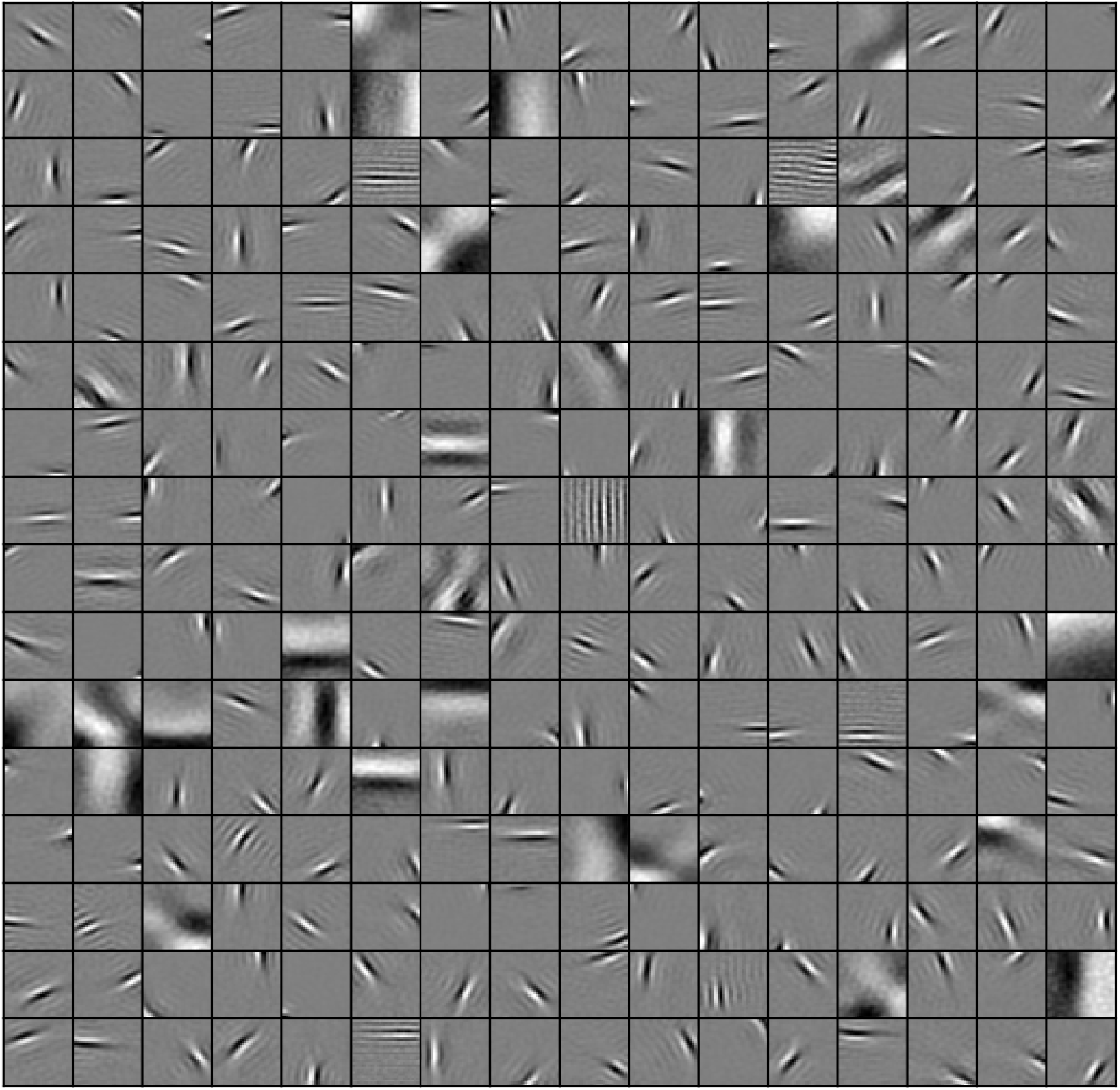
All dictionary elements from a complete sparse coding dictionary learned with stochastic gradient descent and LCA inference on natural image patches that were not prewhitened.

Although these quantities may be available at some location in the neuron, it is not clear how many of them may be available at all of the dendritic synapses during learning. A different criterion for biologically plausible learning rules, and the one we adopted in the main text, is for the rules to be “synaptically local”. Whereas tabulating which neural quantities are available at all dendrites for learning is still an open area of investigation, it is well known that action potentials can backpropagate into the dendritic arbor from the soma [27], so those spike times are likely to be known to all synapses. However, due to electrical compartmentalization throughout the dendrites of typical neurons, it is not clear that the membrane potential at any specific point in the neuron, *e.g.* the soma, is continusously communicated to every synapse in the dendritic tree.

The learning rules for various models require different sets of values to update the dendritic synaptic strengths. For several models, the features required to learn the dendritic synaptic weight for neuron *m* and presynaptic element *j* are listed in Table 1. Here *X*_*j*_ is the *j*th element of the feedforward activity and *X*_*|j*_ are the other elements (not available directly at synapse *mj*). A dendritic synapse may have access to the net filtered input of the feedforward activity, ∑_*j*_ Φ_*mj*_*X*_*j*_, the current membrane potential of its soma, *u*_*m*_, or its own spiking activity and the spiking activity of its neighbors, *a*_*n*_. Finally, gradient descent on the mean-squared reconstruction error requires the dendritic synapse to have access to the sum of the dendritic synapses of all neurons weighted by their spiking activity, ∑_*n*_ Φ_*nj*_*a*_*n*_.

**Table 1:** Features required for learning reconstruction feature weight, Φ_*mj*_, for neuron *m* and presynaptic element *j* for learning rules associated with four models. NL and SL indicate whether the learning rule or quantities is neuron-local (NL) or synaptically-local (SL). Learning rules with a check mark in the “Unwhitened inputs?” row are capable of learning from significantly non-white data.

| Feature | SAILnet<br>[29] | Brendel<br>et al.[5] | MP-SAILnet | mse GD | NL | SL |
| --- | --- | --- | --- | --- | --- | --- |
| $X_j$ | ✓ | ✓ | ✓ | ✓ | ✓ | ✓ |
| $X_{\setminus j}$ | × | × | × | × | ✓ | × |
| $\sum_j \Phi_{mj} X_j$ | × | ✓ | × | × | ✓ | × |
| $a_m$ | ✓ | ✓ | ✓ | ✓ | ✓ | ✓ |
| $u_i$ | × | × | ✓ | × | ✓ | × |
| $\sum_n \Phi_{nj} a_n$ | × | × | × | ✓ | × | × |
| Neuron-local | ✓ | ✓ | ✓ | × |  |  |
| Synaptically local | ✓ | × | × | × |  |  |
| Unwhitened inputs? | × | ✓ | ✓ | ✓ |  |  |

**Figure S4:**
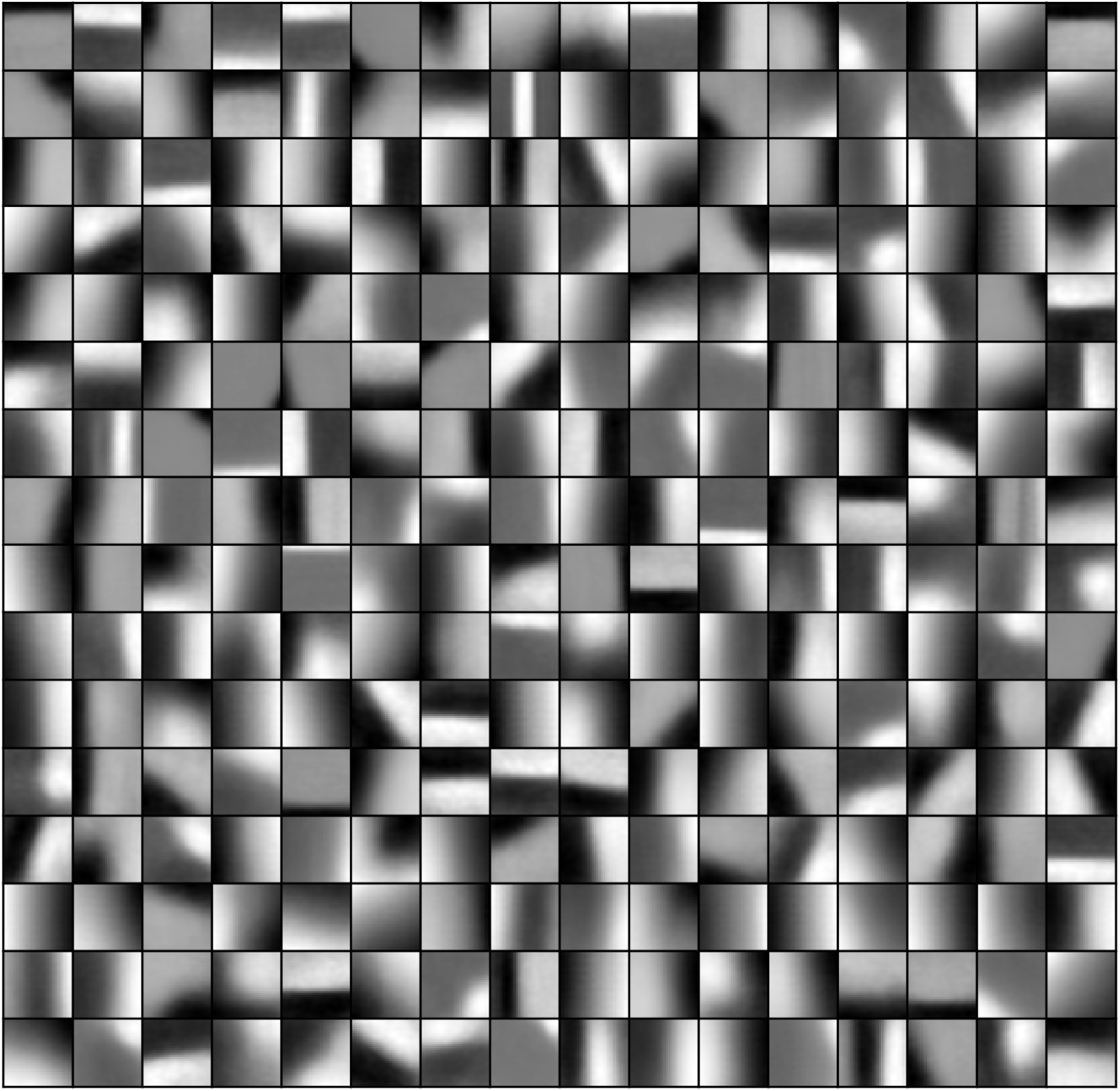
All dictionary elements for a complete SAILnet model trained on natural image patches that were not pre-whitened.

## Acknowledgments

This work was supported in part by the U.S. Army Research Laboratory and the U.S. Army Research Office under Contract No. W911NF-13-1-0390. JAL acknowledges support from the L.D.R.D. Program of Lawrence Berkeley National Laboratory, which is supported by the Office of Science of the U.S. Department of Energy under Contract No. DE-AC02-05CH11231

---

**Figure S5:**
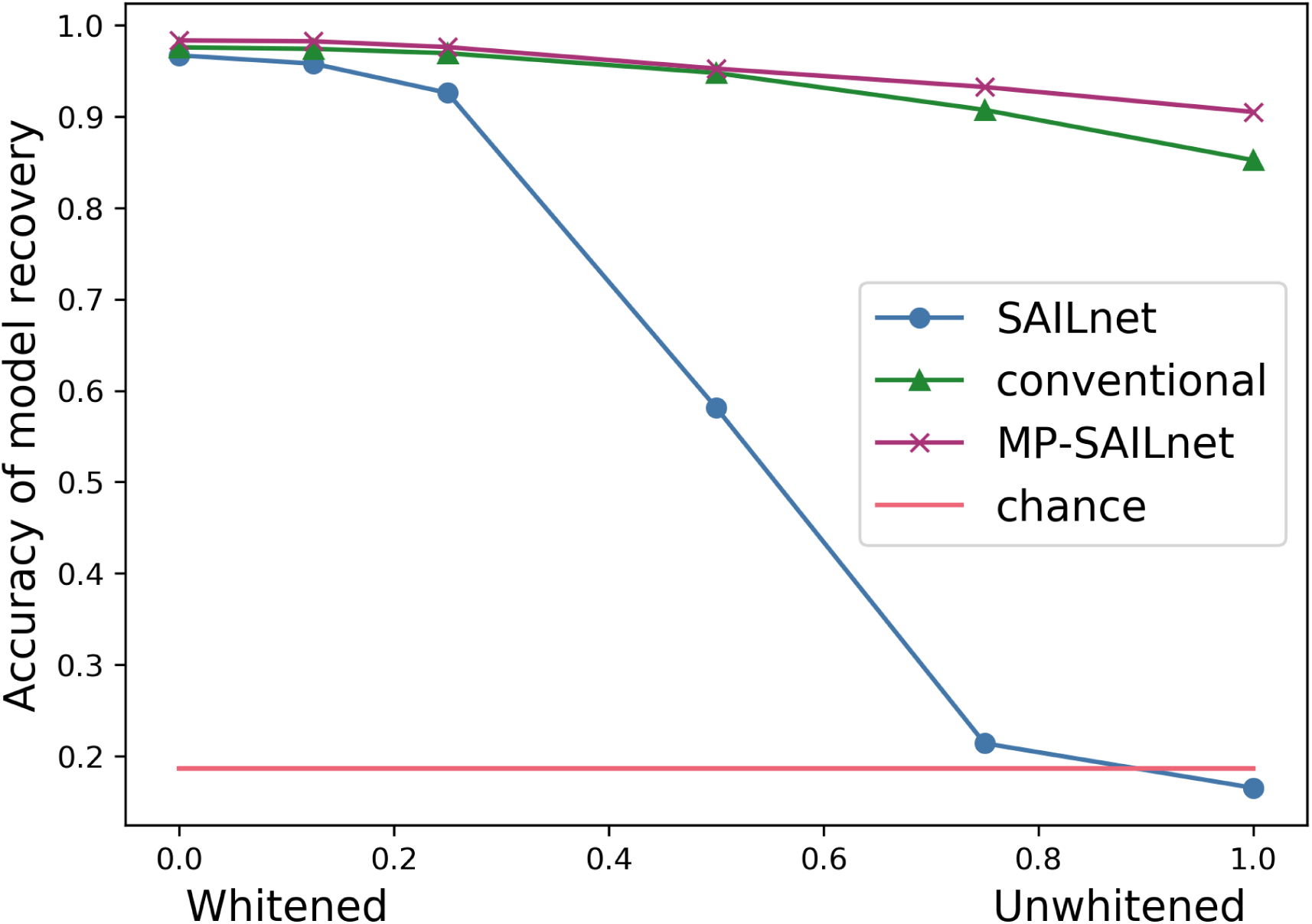
A modified SAILnet with learning rules that depend on post-synaptic membrane potential recovers a sparse model even on unwhitened data. This figure is identical to Fig 1E in the main text, except for the addition of data for our modified SAILnet model (MP-SAILnet).

